# Small extracellular vesicles but not microvesicles from *Opisthorchis viverrini* promote cell proliferation in human cholangiocytes

**DOI:** 10.1101/2023.05.22.540805

**Authors:** Sujittra Chaiyadet, Javier Sotillo, Michael Smout, Martha Cooper, Denise L. Doolan, Ashley Waardenberg, Ramon M. Eichenberger, Matt Field, Paul J. Brindley, Thewarach Laha, Alex Loukas

## Abstract

Chronic infection with *O. viverrini* has been linked to the development of cholangiocarcinoma (CCA), which is a major public health burden in the Lower Mekong River Basin countries, including Thailand, Lao PDR, Vietnam and Cambodia. Despite its importance, the exact mechanisms by which *O. viverrini* promotes CCA are largely unknown. In this study, we characterized different extracellular vesicle populations released by *O. viverrini* (*Ov*EVs) using proteomic and transcriptomic analyses and investigated their potential role in host-parasite interactions. While 120k *Ov*EVs promoted cell proliferation in H69 cells at different concentrations, 15k *Ov*EVs did not produce any effect compared to controls. The proteomic analysis of both populations showed differences in their composition that could contribute to this differential effect. Furthermore, the miRNAs present in 120k EVs were analysed and their potential interactions with human host genes was explored by computational target prediction. Different pathways involved in inflammation, immune response and apoptosis were identified as potentially targeted by the miRNAs present in this population of EVs. This is the first study showing specific roles for different EV populations in the pathogenesis of a parasitic helminth, and more importantly, an important advance towards deciphering the mechanisms used in establishment of opisthorchiasis and liver fluke infection-associated malignancy.

## 1. Introduction

Opisthorchiasis remains a major public health problem in East Asia and Eastern Europe. The main species affecting South East Asia, *Opisthorchis viverrini*, affects near 10 million people, particularly in Thailand and Laos (1). In addition to infection-associated morbidity (including cholangitis, choledocholithiasis and periductal fibrosis), liver fluke infection with *O. viverrini* has been strongly linked to cholangiocarcinoma (CCA), a form of bile duct cancer which has the highest global prevalence in the Northeast region of Thailand (2). Multiple factors are involved in the progression of CCA, including mechanical damage from physical attachment of the liver fluke to the biliary epithelium, chronic immunopathological processes that induce pro-inflammatory cytokines, and the release of parasite-derived excretory/secretory (ES) products (including soluble proteins and extracellular vesicles (EVs)) into the bile ducts that promote cell proliferation (3–6).

These ES products constitute the main players in the crosstalk between the parasite and its host, and blocking this interaction has been shown to eliminate or impair establishment of the worms, reducing cell proliferation and the development of cholangiocarcinoma (3, 6–11). For instance, over 300 proteins have been identified in the ES products from *O. viverrini* (4), one of which is a granulin-like growth factor termed *Ov*-GRN-1, which drives proliferation of biliary epithelial cells. *Ov*-GRN-1-induced cell proliferation can be inhibited with antibodies raised to the recombinant protein (12), and moreover, infection of hamsters with *Ov-grn-1* knock-out flukes results in reduced biliary fibrosis and cholangiocarcinoma compared to hamsters infected with control flukes (8, 13). In addition, we reported that *O. viverrini* secretes small extracellular vesicles (EVs) of 40-100 nm in size that can be internalized by host cells (3), and that blocking internalization of these EVs using antibodies against a member of the tetraspanin protein family significantly reduced the secretion of pro-inflammatory cytokines including IL-6 (3). Furthermore, vaccination of hamsters with small EVs resulted in decreased worm burdens and worm growth retardation as well as reduced egg secretion (9). However, the exact mechanisms by which EVs promote biliary cell proliferation and aids the establishment of *Opisthorchis* infection remain unknown.

EVs can be categorized into different subpopulations based on the origin and size of the vesicles, although, for most cell types and organisms studied, there are no reliable and specific markers for each population (14, 15). Small EVs (usually named exosomes) have an endocytic origin and have a size of 30-150 nm, whereas larger EVs such as microvesicles form by direct budding of the plasma membrane and typically range from 100 to 1000 nm (1 µm) (16). Whereas small vesicles have been described so far from *O. viverrini*, other flukes have been shown to secrete both populations of EVs, including *Fasciola hepatica*, and *Schistosoma mansoni* (17, 18), although their specific roles in parasite-host communication remain unclear.

While the proteomic content of EVs from 17 different helminth species has been well characterized, and several species or class-specific markers have been proposed, the miRNA cargo of EVs has not been characterized for all helminths studied (reviewed in (19)). In schistosomes, parasite-specific miRNAs have been proposed as new diagnostic candidates in infected human subjects (20), and it has been hypothesized that EV miRNAs from the nematode *Nippostrongylus brasiliensis* have anti-inflammatory and immunomodulatory roles (21). Furthermore, *Schistosoma japonicum* EV miR-125b and bantam miRNAs promote macrophage proliferation and TNF-α production (22), and *S. mansoni* miR-10 is implicated in the polarization towards a Th1 response (23) while Sja-miR-71a present in *S. japonicum* egg EVs can suppress liver fibrosis (24). However, no association has been found so far between *O. viverrini* miRNAs and malignancy.

In the present study we show, for the first time, the secretion of both types of EVs by *O. viverrini* and highlight several proteins that are unique for each vesicle type, and that could be used as specific markers for the isolation and characterization of these subpopulations of EVs. Furthermore, we delve into the mechanisms used by the parasite to promote cell proliferation in cholangiocytes via small EVs but not MVs, which highlights the role of EVs in inter-phylum cross-talk and opens new avenues for the treatment of *O. viverrini*-induced cholangiocarcinoma.

## 2. Materials and methods

### 2.1 Animal ethics

Six to eight-week-old male Syrian golden hamsters were infected with *O. viverrini* metacercariae and maintained in the animal house facility at the Faculty of Medicine, Khon Kaen University, Thailand for 8 weeks before being euthanized. Animal experiments were approved by the Animal Ethics Committee of Khon Kaen University (IACUC-KKU 93/2565).

### 2.2 Isolation and purification of extracellular vesicles

Hamsters were necropsied at 8 weeks post-infection and adult worms collected, washed in PBS and cultured in RPMI 1640 containing 1% glucose, 100 units/ml Penicillin, 100 units/ml Streptomycin (Life Technologies, Grand Island, NY) and 1 nM E64 (Thermo scientific, USA) at 37°C, 5% CO_2_ for 7 days. For the isolation and purification of *O. viverrini* EVs (*Ov*EVs), a previously published method was followed (18). Briefly, *O. viverrini* ES products (*Ov*ES) were collected every day, centrifuged at 500 *g* for 10 min to remove eggs and large debris, and subsequently centrifuged at 2,000 *g* for 30 min, 4,000 *g* for 30 min and 15,000 *g* for 45 min to remove smaller debris and MVs. MVs were washed twice with PBS, centrifuged at 12,000 *g* and stored at -80°C until use. Following removal of MVs, supernatant was concentrated using a 10 kDa cut-off Amicon filter (Merck Millipore, USA) and ultracentrifuged at 120,000 *g* for 3 hours to pellet smaller (120k) vesicles. The pellet was resuspended in 70 µl of PBS, laid on a discontinuous gradient (40%, 20%, 10%, 5%) built with OptiPrepTM Density Gradient (Millipore Sigma, USA) as described previously (25) and centrifuged for 18 h at 4°C. EVs isolated from grapes (termed “grape Evs”) were isolated from *Vitis vinifera* Thompson seedless grapes as described elsewhere (21). The size and concentrations of all Evs were analysed using a qNano instrument (Izon Science, New Zealand) and protein content was determined using a BCA kit (Bio-Rad).

### 2.3 Tunable Resistive Pulse Sensing analysis of extracellular vesicles

Tunable resistive pulse sensing (TRPS) was employed using a qNano system (Izon Science) to measure the particle concentration and size distribution of Evs. Briefly, a nanopore NP150 or NP400 (for 12k and 15k Evs, respectively) was used, and pressure and voltage values were set to optimize the signal to ensure high sensitivity. All samples were diluted (1:10 for *Ov*EVs and 1:20 for grapesomes) before applying to the nanopore, and CP100 carboxylated polystyrene calibration particles (Izon Science) were used for calibration. Size distribution and concentration of particles were analyzed using the software provided by Izon (version 3.2).

### 2.4 Proteomic analysis of 15k OvEVs

EV-surface exposed peptides were released by trypsin hydrolysis following the methodology described by Chaiyadet and co-workers (26). Briefly, EVs were treated with trypsin (1 µg/µl) for 15 min at 37°C to cleave surface-exposed proteins, centrifuged at 15,000 *g* for 1 hour at 4°C, and supernatant containing the surface peptides collected. Pellet containing “shaved” EVs was resuspended in water, sonicated and released EV cargo content was recovered from the supernatant after centrifugation at 15,000 *g* for 1 hour at 4°C. Finally, the pellet was solubilized in 1% Triton X-100/2% sodium dodecyl sulphate (SDS) at 37°C for 15 min to recover membrane-associated proteins. For the proteomic analysis, cargo and membrane-associated proteins were electrophoresed on a 10% SDS-PAGE gel and in-gel digestion was performed as described previously with minor modifications (27).

All samples (trypsin-liberated, cargo and membrane peptides) were injected into an Eksigent nanoLC 415 system using an Eksigent Trap-column (C18-CL, 3 μm, 120 Å, 350 μm x 0.5 mm) for the pre-concentration step followed by separation in a 15 cm long Eksigent column (C18-CL particle size 3 μm, 120 Å, 75 μm ID) using a linear gradient of 3-40% solvent B (100 acetonitrile/0.1% formic acid [aq]) in solvent A (0.1% formic acid [aq]) for 45 min followed by 40-80 % solvent B in 5 min. A flow rate of 300 nl/min was used for all experiments. Eluates from the RP-HPLC column were directly introduced into the PicoView ESI ionisation source of a TripleTOF 6600 MS/MS System (AB Sciex) operated in positive ion electrospray mode. All analyses were performed using Information Dependent Acquisition. Analyst Software 1.7.1 (Applied Biosystems) was used for data analysis. Briefly, the acquisition protocol consisted of the use of an Enhanced Mass Spectrum scan with 15 sec exclusion time and 100 ppm mass tolerance. A cycle time of 1800 ms was used to acquire full scan TOF-MS data over the mass range 400–1250 m/z and product ion scans over the mass range of 100–1500 m/z for up to 30 of the most abundant ions with a relative intensity above 150 and a charge state of +2 − +5. Full product ion spectra for each of the selected precursors were then used for subsequent database searches.

Peak lists obtained from MS/MS spectra were identified using X!Tandem version X! Tandem Vengeance (2015.12.15.2) (28), MS-GF+ version Release (v2018.04.09) (29) and Tide (30). The search was conducted using SearchGUI version 3.3.15 (31). Protein identification was conducted against a concatenated target/decoy version of the *O. viverrini* genome and the common repository of adventitious proteins (cRAP database) (10,876 (target) sequences). The decoy sequences were created by reversing the target sequences in SearchGUI. The identification settings were as follows: Trypsin (specific), with a maximum of 2 missed cleavages, 20.0 ppm as MS1 and 0.2 Da as MS2 tolerances; fixed modifications: Carbamidomethylation of C (+57.021464 Da), variable modifications: Deamidation of N (+0.984016 Da), Deamidation of Q (+0.984016 Da), Oxidation of M (+15.994915 Da), fixed modifications during refinement procedure: Carbamidomethylation of C (+57.021464 Da), variable modifications during refinement procedure: Acetylation of protein N-term (+42.010565 Da), Pyrolidone from E (--18.010565 Da), Pyrolidone from Q (--17.026549 Da), Pyrolidone from carbamidomethylated C (--17.026549 Da).

Peptides and proteins were inferred from the spectrum identification results using PeptideShaker version 1.16.40 (32). Peptide Spectrum Matches (PSMs), peptides and proteins were validated at a 1.0% False Discovery Rate (FDR) estimated using the decoy hit distribution. The mass spectrometry data along with the identification results have been deposited in ProteomeXchange Consortium (33) via the PRIDE partner repository with the dataset identifier PXD020356 and doi:10.6019/PXD020356. During the review process, the data can be accessed with the following credentials upon login to the PRIDE website (http://www.ebi.ac.uk/pride/archive/login): Username, Password: [fL0hevyQ].

Blast2GO software (34) was employed for the Gene Ontology (GO) analysis. Only children GO terms were used for subsequent analysis to avoid redundancy in GO terms. Protein family (Pfam) analysis was performed using the gathering bit score (--cut_ga) threshold using HMMER v3.

### 2.4 mRNA and miRNA isolation

mRNA and miRNA of *Ov*EVs were obtained from different batches of worms as described previously (21) with sequencing performed on four mRNA and two miRNA biological replicates of the 120k EVs. Briefly, total RNA and miRNA were extracted using the mirVanaTM miRNA Isolation Kit (ThermoFisher) and stored at −80°C until analysed.

### 2.5 RNA sequencing and transcript annotation

The RNA quality, yield and size of total and small RNAs were analysed using capillary electrophoresis (Agilent 2100 Bioanalyzer, Agilent Technologies, Santa Clara, CA, USA). The TruSeq Small RNA-seq preparation kit (Illumina) was used for miRNA sequencing according to the manufacturer’s instructions. For mRNA-seq, ribosomal RNA was removed from samples, and mRNA was prepared for sequencing using an Illumina TruSeq stranded mRNA seq library preparation kit according to the manufacturer’s instructions. RNAseq was performed on a NextSeq 500 (Illumina, 75-bp PE mid-output run, approximately 30M reads per sample). Quality control, library preparation and sequencing were performed at the Ramaciotti Centre for Genomics at the University of New South Wales, Sydney.

High-throughput RNA-seq data were aligned to the *O. viverrini* reference genome model (WormBase WS255; http://parasite.wormbase.org; (35)) using the STAR transcriptome aligner (36). Differentially expressed genes were identified using consensusDE (37). Prior to downstream analysis, rRNA-like sequences were removed from the metatranscriptomic dataset using riboPicker-0.4.3 (http://ribopicker.sourceforge.net; (38)). BLASTn algorithm (39) was used to compare the non-redundant mRNA dataset for *O. viverrini* EVs to the nucleotide sequence collection (40) from NCBI (www.ncbi.nlm.nih.gov) to identify putative homologues in a range of other organisms (cutoff: <1E-03). Corresponding hits homologous to the murine host, with a transcriptional alignment coverage <95% (based on the effective transcript length divided by length of the gene), and with an expression level <10 fragments per kilobase of exon model per million mapped reads (FPKM) normalized by the length of the gene, were removed from the list. The final list of mRNA transcripts from *O. viverrini* EVs was assigned to protein families (Pfam) and GO categories (Blast2GO).

### 2.6 miRNA analysis and target prediction

The miRDeep2 package (41) was used to identify known and putative novel miRNAs present in both miRNA samples. As there are no *O. viverrini* miRNAs available in miRBase release 21 [36], the miRNAs from the flatworms *Schmidtea mediterranea*, *Echinococcus granulosus* and *Echinococcus multilocularis*, *S. japonicum* and *S. mansoni* were utilized as a training set for the algorithm. Only miRNA sequences commonly identified in all replicates were included for further analyses.

The interaction between human host genes and the identified miRNAs were bioinformatically predicted using three different software programs: MR-microT (42), mirDB (43) and miRANDA (44). miRDB and MRMicroT are web-based algorithms and default settings were used, except that only targets with scores ≥0.7 and ≥60, respectively, were selected. In the case of miRANDA, input 3ʹUTR from the *Homo sapiens* GRCm38.p4 assembly was retrieved from the Ensembl database release 100. The software was run with strict 5ʹ seed pairing, energy threshold of −20 kcal/mol and default settings for gap open and gap extend penalties. Interacting hits were filtered by conservative cut-off values for pairing score (>155) and matches (>80%). For a more robust target prediction, only targets commonly predicted by all three software for a single miRNA were further analysed by the Panther classification system (http://pantherdb.org/) using pathway classification (45).

### 2.7 Mammalian cell culture

The nonmalignant cholangiocyte cell line H69 is a SV40-transformed human bile duct epithelial cell line derived from human liver, kindly provided by Dr. Gregory J. Gores, Mayo Clinic, Rochester, Minnesota. H69 cells (46) were maintained in T75 cm^2^ vented flasks (Corning) as monolayers as described (47) with minor modifications. Cells were maintained with regular splitting using 1x TrypLE express trypsin (Gibco) every 2–5 days in complete media (Gibco): growth factor-supplemented DMEM/F12 with high glucose media with 10% FCS, 2× antibiotic/antimycotic, 25 μg/mL adenine, 5 μg/mL insulin, 1 μg/mL epinephrine, 8.3 μg/mL holo-transferrin, 0.62 μg/mL hydrocortisone, 13.6 ng/mL T3, and 10 ng/mL EGF. Low nutrient media for cell proliferation assays was 5% complete media, i.e., 0.5% FCS and 1/20 of the growth factor concentrations listed above for complete media. The identity (human derived) of the cell line was confirmed with single tandem repeat (STR) analysis in January 2018 (15/15 positive loci across two alleles) and mycoplasma free status was determined at the DNA Diagnostics Centre (DDC)–Medical (U.S.A.), accredited/certified by CAP, ISO/IEC 17025:2005, through ACLASS.

### 2.8 Cell Proliferation Monitoring in Real Time Using xCELLigence

Cells were seeded at 3,000 cells/well in 150 μL of complete media (above) in E-plates (Agilent) and grown overnight while being monitored with an xCELLigence SP system (Agilent), which monitors cellular events in real time by measuring electrical impedance across interdigitated gold microelectrodes integrated into the base of tissue culture plates. As previously described (48), cells were washed three times with PBS prior to addition of 150 μL of low nutrient media (above) and incubated for a minimum of 6 h before further treatment. Treatments were prepared at 8.5× concentration and added to each well in 20 μL, for a final in-well 1× concentration. The xCELLigence system recorded cell indexes at intervals of 1 h for 5–6 days following treatment. Readings for the cell index were normalized prior to treatment, and cell proliferation ratios represent the relative numbers of cells to controls at each timepoint from 6 replicates. Dose response curves for each treatment at day 3 were from two experiments.

### 2.8 Statistical analysis

Comparisons of induction of cell proliferation in response to treatments were accomplished using Two-Way ANOVA test with Holm-Sidak multiple comparison correction, using GraphPad Prism 6.03. Data were expressed as the mean ± standard error of three independent experiments using Graphpad Prism Software Version 6.03 (www.graphpad.com).

## 3. Results

### 3.1 Purification of EVs

Two different populations of *Ov*EVs were isolated and purified from the *Ov*ES products. The 15k *Ov*EVs were isolated and purified by ultracentrifugation, while 120k *Ov*EVs and grape EVs were further purified using Optiprep® gradient and highly pure fractions combined as described previously (21). Particle diameter and concentration was measured using a qNano instrument, protein concentration using the BCA assay and purity was obtained following Webber and Clayton approach (49). While 15k *Ov*EVs had a mean particle diameter of 320±138 nm (mode 246 nm) and a total concentration of 1.06E+09 particles/ml, the 120k *Ov*EVs had a mean particle diameter of 135±32.7 nm (mode 117 nm) and a total concentration of 4.27E+10 particles/ml (Figure 1). Grape EVs had a mean particle size of 143±30.4 nm and a concentration of 2.41E+10 (Figure 1). Purity of vesicles ranged from 4.61E+07 for the 15k *Ov*EVs to 1.82E+09 for 120k *Ov*EVs (Supplementary Table 1).

**Figure 1.**
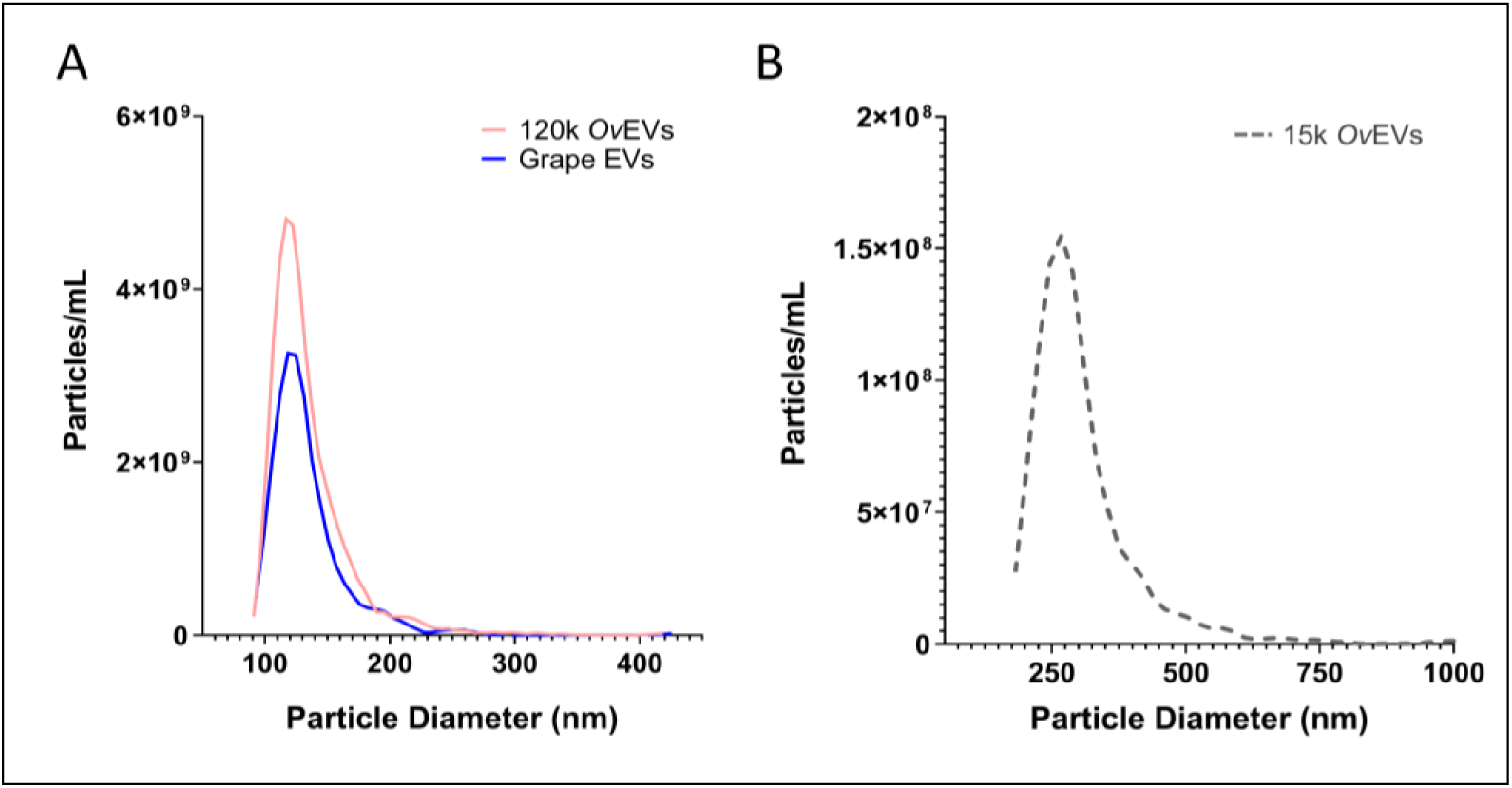
Size and concentration of *Opisthorchis viverrini*-derived 120 k and 15k extracellular vesicles (EVs). Tunable Resistive Pulse Sensing was used to analyse the diameter and concentration of 120k *O. viverrini* EVs as well as grape EVs (A) and 15k *O. viverrini* EVs.

### 3.2 120k OvEVs but not 15k OvEVs promote cholangiocyte proliferation in vitro

To further investigate the role of the different populations of *Ov*EVs in driving proliferation of cholangiocytes, H69 immortalised cholangiocytes were incubated with different concentrations of 120k and 15k *O. viverrini* EVs; grape EVs and media alone were used as controls. 120k EVs significantly promoted cell proliferation from 9.3E+07 up to 7.4E+08 particles/ml (corresponding to 50 ng/ml and 400 ng/ml, respectively) in all concentrations tested (Figure 2A, *P*<0.0001). *O. viverrini* 120k EVs promoted cell proliferation 24 h after incubation with cholangiocytes at different concentrations when compared to controls (Figure 2B, *P*<0.05-0.0001).

**Figure 2.**
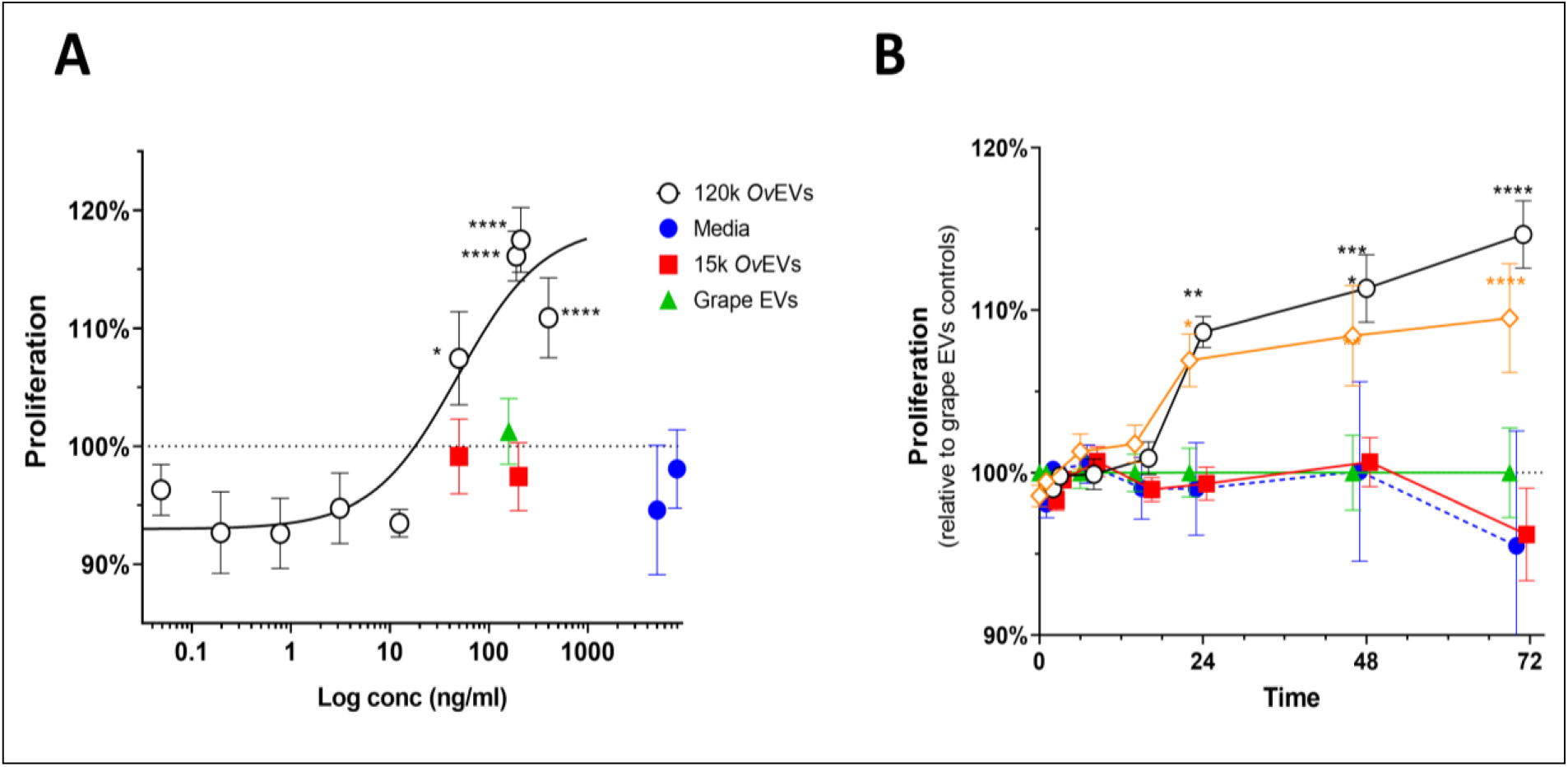
*Opisthorchis viverrini* 120k but not 15k extracellular vesicles (EVs) drive proliferation of human cholangiocytes. (A) *O. viverrini* 120k EVs (open circles) but not 15k EVs (red squares) promoted proliferation of human cholangiocytes at different concentrations when compared to media alone and to grape EVs (blue circles). (40) *O. viverrini* 120k EVs at 3.7E+08 particles/ml (200 ng/ml) (open circles) and 7.4E+08 particles/ml (400 ng/ml) (orange diamond) promoted proliferation of human cholangiocytes significantly after 24 h of incubation.

### 3.3 Proteomic profile of 15k EVs secreted by O. viverrini

The proteomic content of the 15k *Ov*EVs was analysed by mass spectrometry following established methods designed to localise the proteins within the different fractions of the EVs (17, 26). A total of 718 unique proteins were identified with 2 or more unique peptides, including 334 as trypsin-liberated (or “shaved”) proteins, 352 as membrane proteins and 648 as cargo proteins (Supplementary Tables 2-4). A comparison of the results obtained in this analysis was performed against the proteomic data obtained by Chaiyadet et al. (26), since both analyses were performed following the same methodology and using the same mass spectrometers. Interestingly, the cargo from the 15k and 120k *Ov*EVs have the highest number of unique proteins (155 and 49, respectively), while 76 proteins were common to both populations of *Ov*EVs (Figure 2). Furthermore, 46 and 21 proteins are uniquely present in the shaved fractions analysed from the 15k and the 120k population, respectively. (Figure 3, Tables 1, 2).

**Figure 3.**
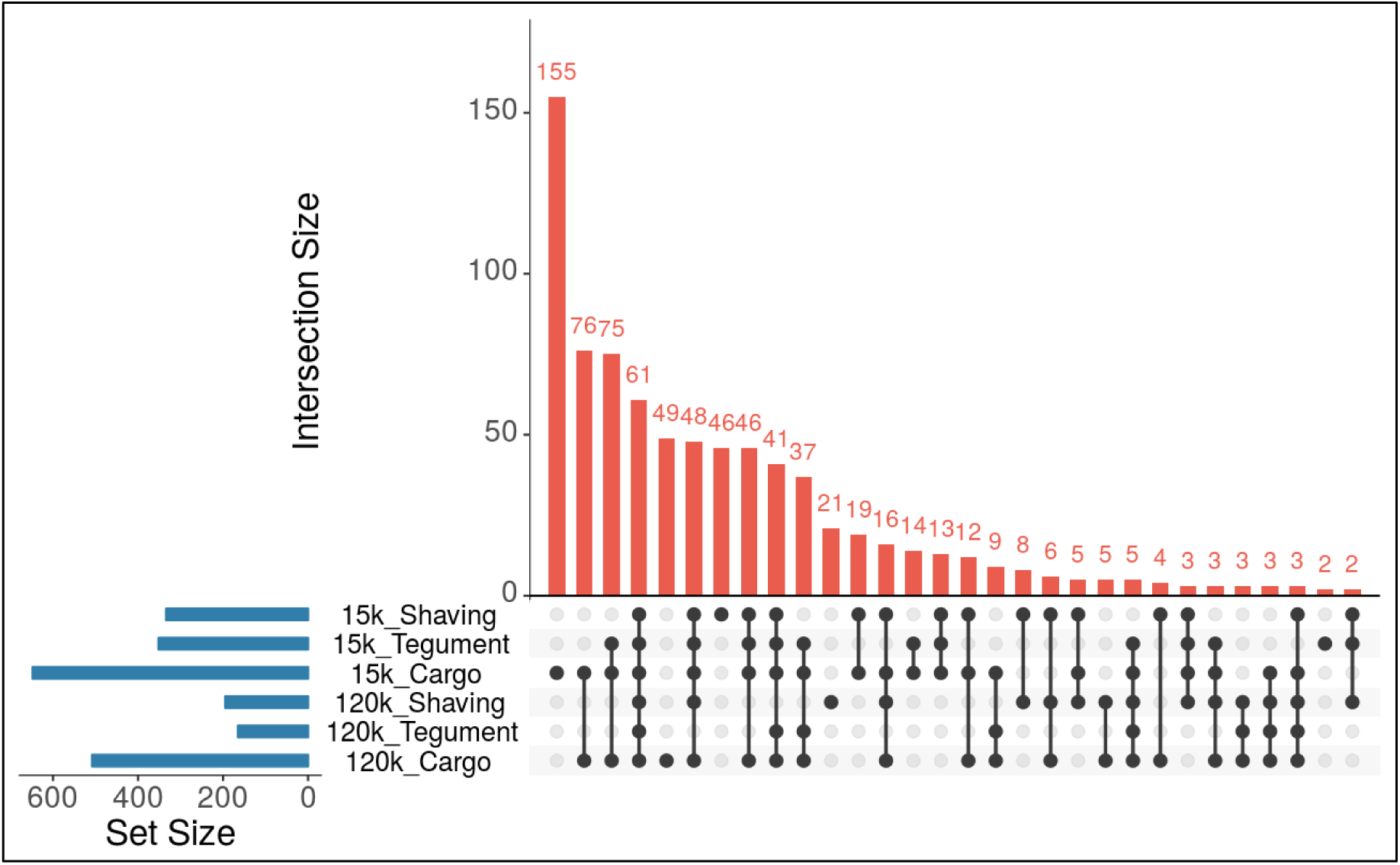
Proteomic identifications of the 120k and 15k populations of *O. viverrini*-derived extracellular vesicles (*Ov*EVs). UpSetR analysis visualising intersections between the proteins identified in the different compartments analysed from the 120k and 15k populations of *O. viverrini*-derived EVs.

**Table 1.** Unique proteins from 120k *Ov*EVs shaved sample.

| Protein name | Description | Pfam family/domain | N° unique peptides |
| --- | --- | --- | --- |
| OON13804.1 | Cell division cycle and apoptosis regulator protein 1 | SAP Family | 6 |
| OON21748.1 | Transcription elongation regulator 1 | FF Family / WW Domain | 5 |
| OON22257.1 | TRPM8 channel-associated factor 2 | Peptidase_M60 Domain | 4 |
| OON21215.1 | Hsp90 protein | HSP90 Family | 4 |
| OON20288.1 | Immunoglobulin domain protein | Ig_3 Domain | 3 |
| OON19373.1 | Oxidoreductase, 2OG-Fe(II) oxygenase family protein | - | 3 |
| OON16932.1 | Putative chaperone protein DnaK | HSP70 Family | 3 |
| OON23969.1 | Ribosomal protein L19e | Ribosomal_L19e Family | 2 |
| OON21776.1 | Large subunit ribosomal protein L12.e | Ribosomal_L11_N Domain | 2 |
| OON20086.1 | Hypothetical protein X801_04039, partial | Vinculin Family | 2 |
| OON19823.1 | High mobility group protein DSP1 | HMG_box_2 Domain | 2 |
| OON18290.1 | Elongation factor 1-beta | EF-1_beta_acid Domain | 2 |
| OON17975.1 | Protein hu-li tai shao | Aldolase_II Domain | 2 |
| OON17798.1 | Putative maleylacetoacetate isomerase | GST_N_3 Domain | 2 |
| OON16907.1 | Neuroblast differentiation-associated protein AHNAK | - | 2 |
| OON16165.1 | Linker histone H1 and H5 family protein | Linker_histone Domain | 2 |
| OON16018.1 | Histone H3 | Histone Domain | 2 |
| OON15665.1 | Carbohydrate phosphorylase | Phosphorylase Family | 2 |
| OON15027.1 | D-aspartate oxidase | DAO Domain | 2 |
| OON13880.1 | Putative cystathionine beta-synthase | PALP Family | 2 |
| OON13684.1 | Cell division protein | Septin Domain | 2 |

**Table 2.**
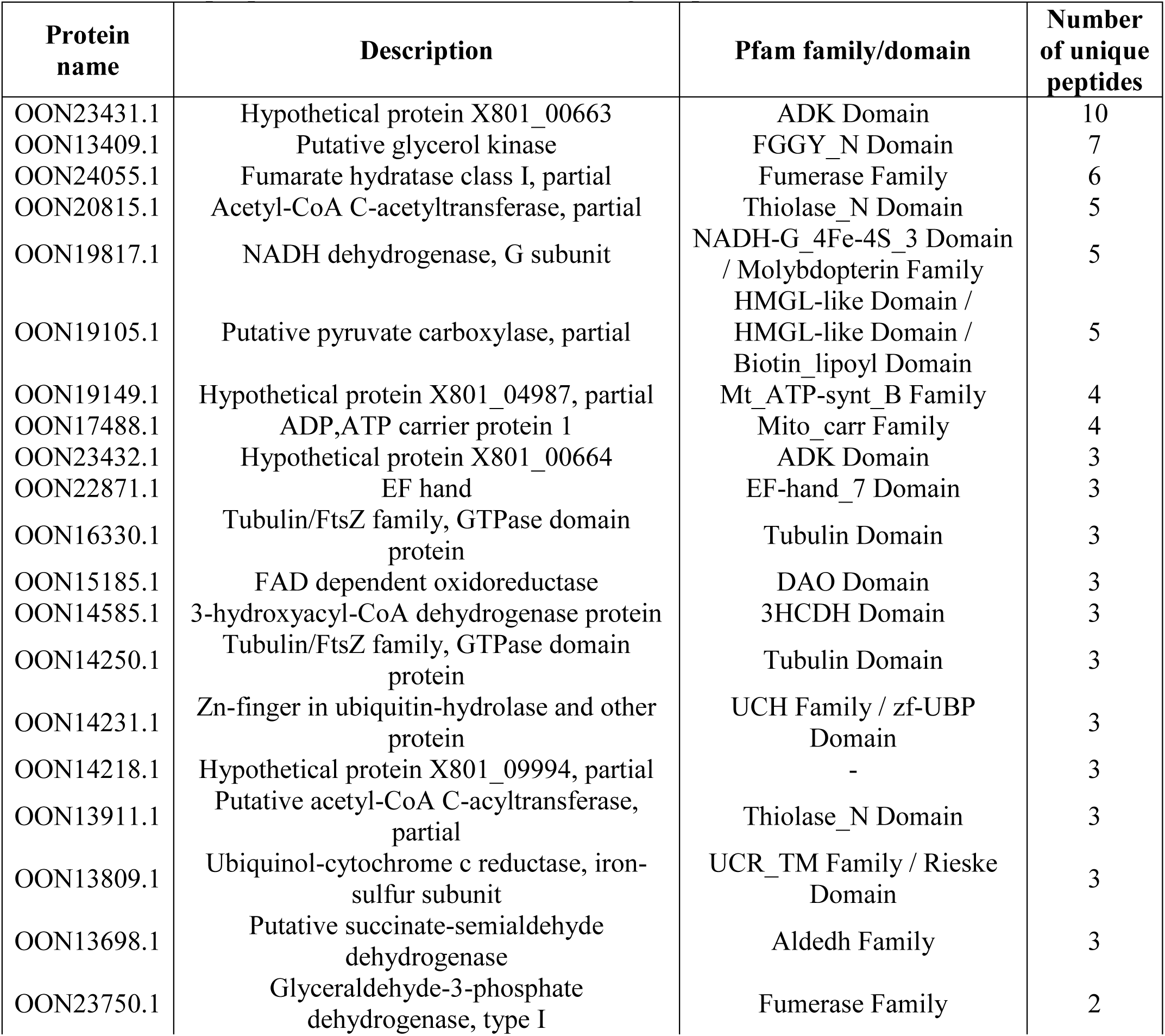
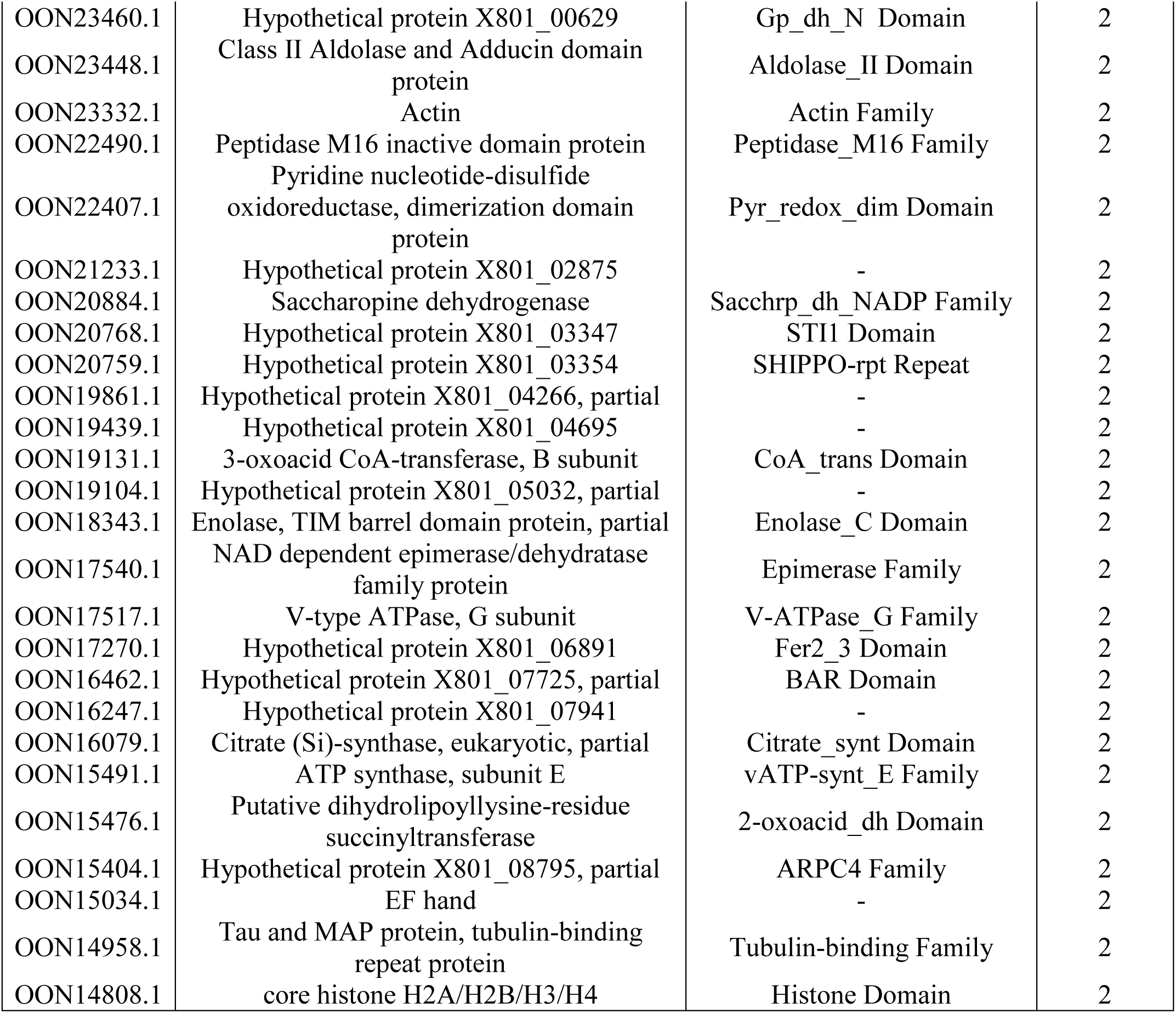
Unique proteins from 15k *Ov*EVs shaving sample.

### 3.4 Characterisation of the genomic content and in-silico target prediction of miRNAs present in 120k OvEVs

By sequencing and screening biological duplicates for mRNAs and miRNAs in the cargo of 120k *Ov*EVs, a total of 2,478 full-length mRNA transcripts mapping to *O. viverrini* gene models were common to both replicates (Supplementary File 1). The identified hits were subjected to a Pfam and GO analysis. The most represented biological processes based on the nodescore provided by Blast2GO were signal transduction (GO:0007165) and transport (GO:0006810), with 195 and 227 sequences matching these processes respectively (Supplementary Figure 1). Similarly, metal ion binding (GO:0046872) and ATP binding (GO:0005524) were the molecular functions with the highest nodescore. Interestingly, proteins encoded by these mRNAs contained at least 1,640 domains based on a Pfam analysis, with Reverse transcriptase (PF00078) and Protein kinase (PF00069) being the most represented (Supplementary Table 5). Similarly, 64 miRNAs were common to both biological replicates, 26 of which have close homologues in other trematodes (Supplementary Table 6).

Potential interactions of *O. viverrini* 120k EV miRNAs with human host genes was explored by computational target prediction. To obtain more robust target predictions, three different target prediction software programs were used and only common targets predicted by all three software for a particular miRNA were taken into consideration for further analysis. From the total 64 miRNAs identified, 50 had targets commonly predicted by all three software (Supplementary Table 7), and, from these, 30 had targets that could be mapped to biological pathways by PantherDB. A total of 85 different pathways could be mapped by the predicted miRNA targets, with the Gonadotropin-releasing hormone receptor pathway being the pathway mapped to the greatest number of genes (22 predicted targets from 18 different miRNAs). Interestingly, genes belonging to different pathways associated with the immune system such as the Inflammation mediated by chemokine and cytokine signaling pathway (P00031), T cell activation (P00053), B cell activation (P00010), Interleukin signaling pathway (P00036) and TGF-beta signaling pathway (P00052) were targeted by *O. viverrini* 120k EV miRNAs (Figure 4, Supplementary Table 7). Genes from different signaling pathways (i.e. Wnt signalling pathway (P00057), the Integrin signalling pathway (P00034) or the PDGF signaling pathway (P00047)) were also predicted to be targeted by *O. viverrini* 120k EV miRNAs. Other pathways targeted by these miRNAs were the Angiogenesis (P00005) pathway, the Apoptosis signaling pathway (P00006) and the VEGF signaling pathway (P00056) (Figure 4, Supplementary Table 7).

**Figure 4.**
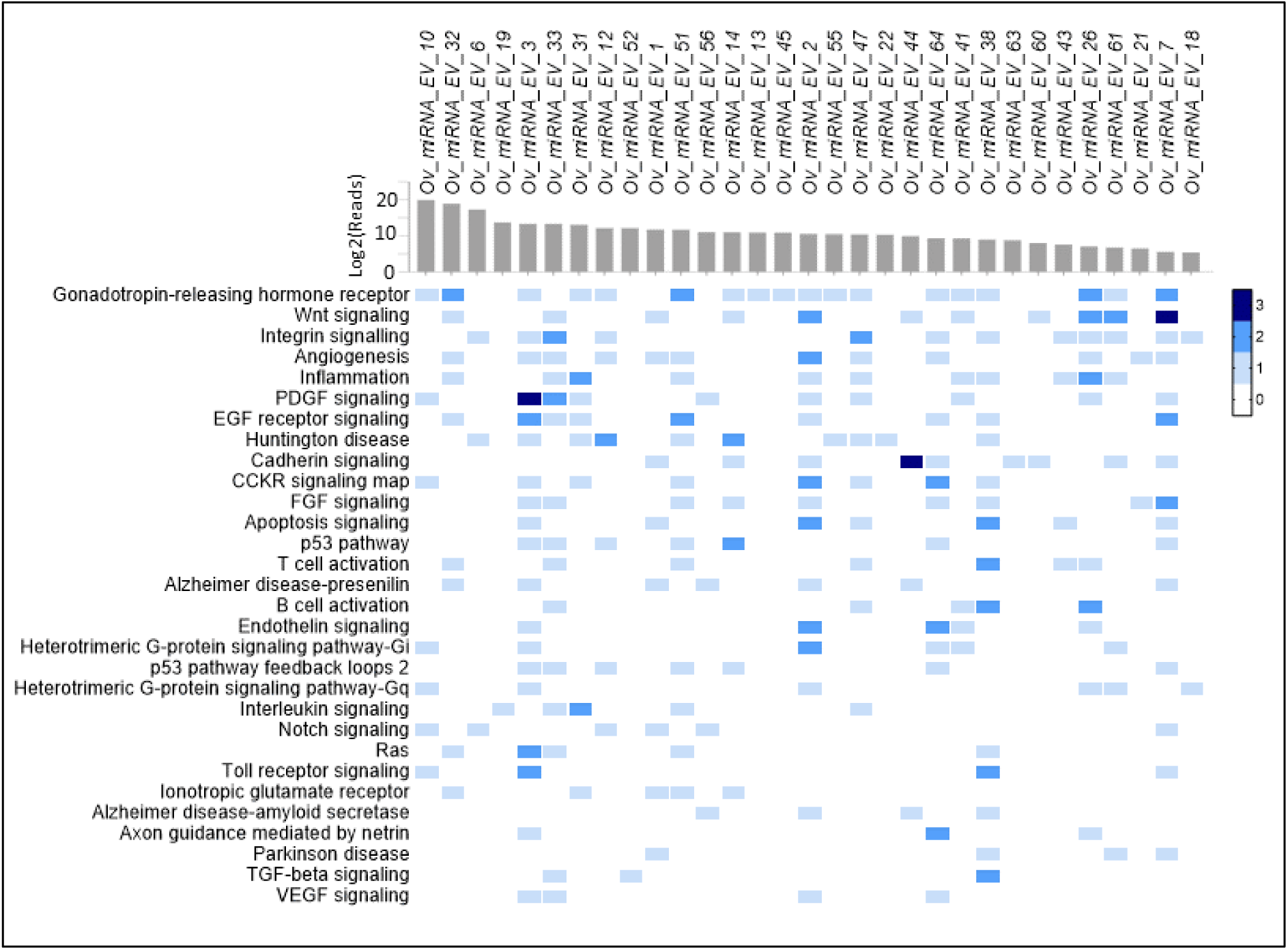
Prediction of *O. viverrini* 120k extracellular vesicle (EV) miRNA target interactions to human host genes. Individual targeted host genes were categorized by PantherDB pathway analysis (heatmap corresponds to individual targeted genes commonly identified by three different target prediction software in the human host). Top axis shows the 30 identified miRNAs in *O. viverrini* EVs containing at least one gene involved in a biological pathway and their abundances (Log2 average mean read counts from two biological replicates).

## Discussion

Discrete types of EVs can be produced by most types of cells, including exosome-like EVs (30-150 nm in diameter) that derive from the endosomal pathway and are formed by inward budding of the multivesicular body membrane, or MVs, which are larger in diameter (100-1,000 nm) and bud directly from the plasma membrane (50). Species of parasitic helminths have been shown to produce both types of EVs, including the human blood fluke, *S. mansoni* and the livestock liver fluke, *Fasciola hepatica* (17, 18). We now report that *O. viverrini* produces at least 20 times more 120k *Ov*EVs than 15k *Ov*EVs. This contrasts with what was observed for *S. mansoni*, which produced more 15k *Sm*EVs than 120k *Sm*EVs (18). However, those studies were performed using only one biological replicate of EVs, and additional replicates should be analysed to confirm these data, as well as to validate it using additional parameters for normalization as recommended in the recently published guidelines for the study of extracellular vesicles from parasitic helminths (15). Our study found that 15k *Ov*EVs are significantly larger in diameter than 120k *Ov*EVs.

Infection with *O. viverrini* remains a major public health concern in liver fluke endemic regions given the risk of CCA or bile duct cancer (PMID: 34504109) . Despite notable efforts to determine the molecules involved in the promotion of cell proliferation and tumorigenesis, the exact mechanisms remain only partially defined (6). For instance, single molecules such as *Ov*-GRN-1 have been strongly linked to cell proliferation, although, it has also recently been demonstrated that more complex structures such as EVs can also induce the release of cytokines involved in fibrosis and tumour progression (3). Indeed, blocking internalization of EVs by host target cells using specific antibodies can significantly reduce cholangiocyte proliferation and release of proinflammatory IL-6 (3). Notably, antibodies targeting the tetraspanin protein, *Ov*-TSP-1 (OON13593.1 in this study), which is found on the surface of both 120k and 15k *Ov*EVs, was able to block EV uptake by host cells. Tetraspanins have been shown to be key molecules for both the release of EVs by donor cells and the internalization of EVs by target cells (51); however, despite their utility as markers to differentiate EV subpopulations in other organisms (52), we have not yet detected members of this family of proteins that are uniquely present in either population. This is likely due to the physiological nature of the trematode tegument, which is enriched in tetraspanins. Chaiyadet et al. also studied the importance of other tetraspanins in the secretion of EVs (26). Using RNAi they showed that silencing the expression of two different genes encoding tetraspanins (*Ov*-TSP2 and *Ov*-TSP3; OON16870.1 and OON14450.1, respectively in our study), resulted in significant reductions in 120k *Ov*EV production (26). Studies analysing the impact of RNAi on the release of 15k *Ov*EVs should be performed in the future.

Although we did not detect tetraspanins that were unique to either type of *Ov*EV and therefore of value as specific vesicle markers, trypsinization of the EV membranes revealed 21 and 46 proteins specifically present on the 120k or 15k *Ov*EVs, respectively. Since these molecules have extra-vesicular domains and peptides that can be potentially targeted by antibodies, they could be used to discriminate between both types of EVs, even in *in vivo* experiments requiring labelling of native structures. Among these molecules we identified members of the heat shock protein 90 and 70 families, peptidases and immunoglobulin-like proteins in the case of 120k *Ov*EVs, and kinases, hydratases, acetyltransferase and EF-hand domain containing proteins in the case of 15k *Ov*EVs. Interestingly, a recent report analysing all proteins present in EVs from trematodes, nematodes and cestodes found that proteins belonging to the heat-shock protein 70 and EF-hand protein families were present in EVs from all trematodes studied (19), although this report did not differ between EV types. Furthermore, this study found that proteins belonging to the Tubulin and Tubulin_C families were more represented in datasets of EVs secreted by liver flukes compared to other helminths (19). Clearly, both populations of *Ov*EVs contain different types of proteins, which might exert diverse functions in recipient cells. Indeed, it is tempting to speculate that the different proteins present on the surface of 120k and 15k *Ov*EVs might target different populations of host cells, contributing to cell-specific effects in the host.

In addition to proteins, genetic material can also be transported by EVs as a means of cell-to-cell communication, modifying the translational profile of the recipient cell and exerting different functions (19, 53, 54). In helminths, the effects of miRNAs on host gene expression has primarily been investigated in relation to immune modulation and parasite survival (53, 55, 56). Despite their important roles, the miRNA content of only four trematode species has been investigated so far, including *Dicrocoelium dendriticum*, *F. hepatica*, *S. mansoni* and *S. japonicum* (57–63). miRNAs mir-71, mir-10, mir-190, let-7 and mir-2 were present in all four trematode species with relative abundance (19). In our study, miRNAs belonging to the mir-71 and mir-10 families (Ov_miRNA_EV_32 and Ov_miRNA_EV_6, respectively) were among the most abundant in 120k *Ov*EVs. mir-71 has also been shown to be abundantly expressed by *Clonorchis sinensis*, and has been suggested to have a role in parasite survival and metabolism inside the host (64). Furthermore, other miRNAs from the let-7 and mir-2 families were also identified in this sample. Let-7 has been suggested as a potential biomarker for cestode infections (65, 66), while miR-2 has been suggested to regulate growth, development and parasite-host interaction during the migration within the definitive host (67). We used three different target prediction software programs to bioinformatically predict potential host genes that could be the target of the identified miRNAs. Furthermore, to obtain more robust target predictions, only common targets predicted by all three software for a particular miRNA were taken into consideration for further analysis. We also grouped the identified targets into the pathways in which they participate to obtain a broader picture of the effects of miRNAs in the host. Of note, the Gonadotropin-releasing hormone receptor pathway was the pathway to which the greatest number of genes mapped (22 predicted targets from 18 different miRNAs). Interestingly, this pathway has been strongly linked to CCA and other cancers such as pancreatic cancer (68, 69). Although the knockdown of Gonadotropin-releasing hormone decreased cholangiocyte proliferation and fibrosis, this hormone has also been shown to inhibit cell proliferation in other carcinomas including breast, pancreas and liver (70). Furthermore, genes involved in other pathways such as the Angiogenesis (P00005) pathway, the Apoptosis signaling pathway (P00006) and the VEGF signaling pathway (P00056), all implicated in the development of CCA, can also be targeted by miRNAs present in the 120k *Ov*EVs (71, 72). Interestingly, other pathways involved in immunomodulation can also be targeted by miRNAs present in the 120k *Ov*EVs, which might contribute to the formation of an inflammatory microenvironment that favours the development of fibrosis and CCA. Ideally, the roles of the miRNAs involved in the different pathways found in this study should be individually validated in future studies.

In this work we have delved deeper into the responses induced by different populations of fluke EVs in human cholangiocytes with a particular focus on cell proliferation. Furthermore, we have performed a proteomic comparative analysis and the first miRNA analysis of 120k *Ov*EVs to obtain a more accurate picture of the mechanisms used by *O. viverrini* in host-parasite interactions. Finally, the proteomic and miRNA transcriptomic analyses performed will also allow us to identify specific proteins that could be used to discriminate both types of vesicles as well as potential miRNAs implicated in pathogenesis.

## Acknowledgments

This research was supported from a project grant from the National Health and Medical Research Council of Australia (NHMRC), grant identification number APP1085309, the National Cancer Institute, National Institutes of Health, award number R01CA164719, and the Fundamental Fund, Khon Kaen University. AL is supported by a Level Three NHMRC Investigator Grant APP2008450. JS is supported by a Ramon y Cajal fellowship (RYC2021-032443-I) from the Ministerio de Ciencia e Innovacion from Spain.

## Author Contributions

S.C., J.S., A.L., and T.L. conceived the project and in the design of the experiments. S.C., J.S., M.S., R.M.E., M.F. and A.W. performed the experiments and analyzed the results. All authors reviewed the manuscript.

## Supplementary Figures

Figure S1. Gene ontology (GO) analysis of molecular function terms associated to proteins present in adult *O. viverrini*-derived 15k (A) and 120k vesicles (40). Similarity based scatterplots showing the most abundantly represented GO terms ranked by nodescore (Blast2GO). Increasing heatmap score signifies increasing nodescore from Blast2GO, while circle size denotes the frequency of the GO term from the underlying database.

Figure S2. Gene ontology (GO) analysis of biological process terms associated with proteins present in adult *O. viverrini*-derived 15k (A) and 120k vesicles (40). Similarity based scatterplots showing the most abundantly represented GO terms ranked by nodescore (Blast2GO). Increasing heatmap score signifies increasing nodescore from Blast2GO, while circle size denotes the frequency of the GO term from the underlying database

