## Supplementary figures and images for "Small extracellular vesicles but not microvesicles from *Opisthorchis viverrini* promote cell proliferation in human cholangiocytes"

### Figure S1

#### Slide 1
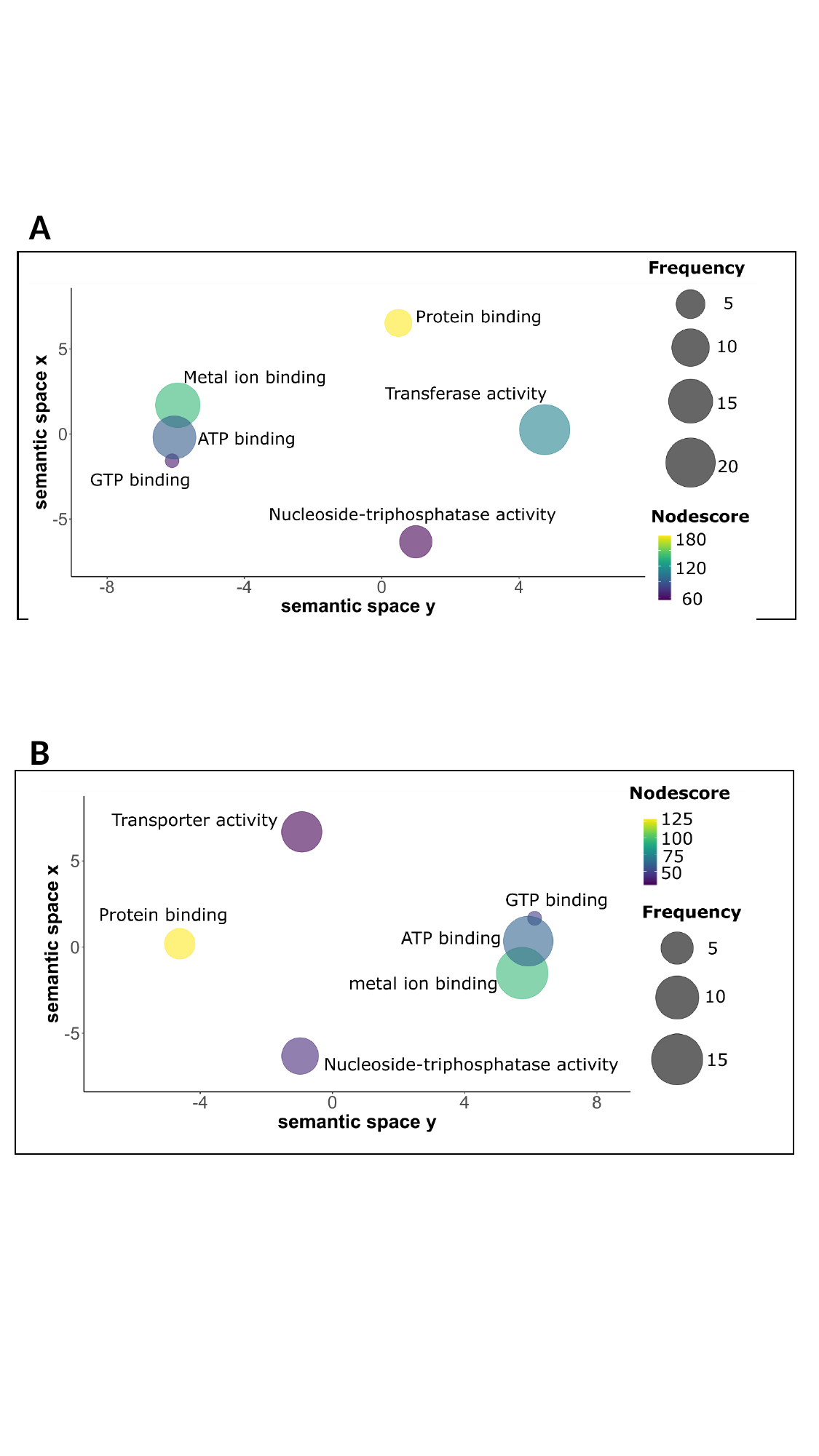

A
B

### Figure S2

#### Slide 1
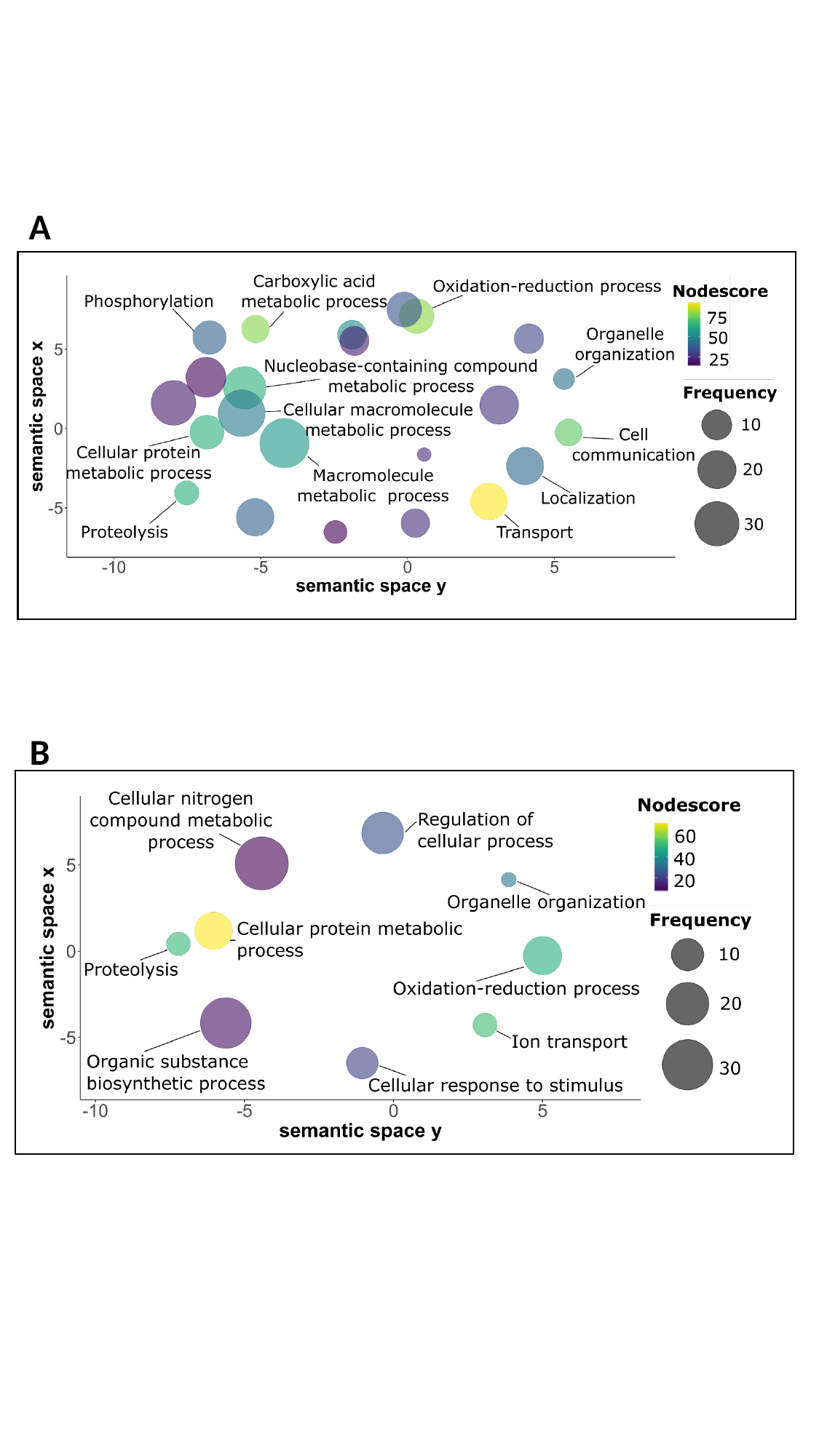

A
B
